# MRI-based dental maturity in newborns reflects prenatal exposures and predicts timing of primary tooth eruption

**DOI:** 10.64898/2026.05.21.726980

**Authors:** Ying Meng, Thomas G. O’Connor, Felicitas B. Bidlack, Scotty A. Simmons, Jin Xiao, Jerod M. Rasmussen

## Abstract

Primary tooth development is shaped by prenatal experience and has consequences for childhood caries and lifelong oral health. However, evidence linking prenatal conditions to dental development has relied largely on postnatal proxies, such as parent-recalled eruption timing, that conflate prenatal programming with postnatal exposures. Here, we directly phenotype developing dentition in vivo using routinely acquired neonatal brain MRI. Using T2-weighted imaging from the HEALthy Brain and Child Development Study, we trained a 3D U-Net on 100 semi-automatic labels and applied the model to 1,433 quality-controlled neonatal scans. Automated post-processing extracted quantitative features of tooth volume, mineralization, and arch geometry. These features were used to predict postmenstrual age at MRI and to derive a bias-corrected tooth age gap (TAG), indexing relative dental maturity at birth. The segmentation model achieved mean cross-validated Dice = 0.94. Dental features predicted postmenstrual age with R^2^ = 0.30 and mean absolute error = 6.7 days, outperforming standard anthropometric measures (R^2^ = 0.22). In adjusted models, TAG varied by infant sex and was associated with gestational age at delivery, maternal pre-pregnancy BMI, and prenatal tobacco use. In infants with longitudinal dental follow-up, greater dental maturity at birth predicted earlier first tooth emergence and more teeth at one year. Automated segmentation and age prediction generalized to an independent cohort. These findings establish neonatal dental MRI phenotyping as an objective, scalable index of dental maturation and a potential readout of prenatal influences on a developmental system relevant to lifelong oral health.

**Significance Statement:** Primary tooth development begins before birth and is shaped by intrauterine experience, with consequences for eruption timing, enamel quality, childhood caries, and lifelong oral health. Yet newborn dentition has been difficult to study because unerupted primary teeth are not routinely measurable at birth, forcing reliance on postnatal proxies that mix prenatal influences with later exposures. We show that standard newborn brain MRI can be repurposed to quantify dental maturity before postnatal exposures accumulate. Applied to 1,433 newborns, this open-source pipeline provides spatially localized measures of unerupted tooth development and links neonatal dental maturity to clinical, environmental, and eruption outcomes. These findings establish neonatal dental MRI as a scalable tool for studying how intrauterine experience becomes biologically embedded in dental development.

## 1. Introduction

Dental caries is the most prevalent chronic disease of childhood, affecting over half of children aged 6–9 years (1). Two contributors to caries susceptibility, primary tooth eruption timing and enamel quality, can be traced back to pre/perinatal development, a period when primary teeth are actively forming but remain hidden from clinical observation (2-4). Primary teeth begin development during the fifth to sixth week of gestation, and crown mineralization begins by the second trimester and continues into early life (5). Because enamel does not remodel after formation, primary teeth can preserve a durable biological record of the intrauterine environment (6). Thus, measuring primary tooth development at birth may help identify early variation in dental traits relevant to childhood caries risk.

A growing body of evidence links prenatal and perinatal exposures and conditions to variation in primary dentition. For example, maternal smoking has been associated with accelerated eruption (3), preterm and early birth with delayed eruption and increased enamel defects (7, 8),and maternal nutritional status with enamel hypoplasia and hypomineralization (8, 9). Further,several of these exposures/conditions are also linked to elevated early childhood caries risk (10). Yet this literature remains heterogeneous and limited by measurement. Specifically, existing studies rely on parent-recalled eruption timing or post-eruption clinical assessment of enamel,both of which are measured after postnatal exposures have begun to accumulate and neither of which directly characterizes unerupted dental development at birth.

Standard newborn brain MRI protocols offer an unexploited opportunity to address this gap; they routinely acquire whole-head volumes that include the developing dentition within the maxilla and mandible. On T2-weighted sequences, the water-rich cellular layers that will form enamel and dentin appear hyperintense, while the mineralized matrix these cells deposit appears hypointense. The resulting natural tissue contrast is well-suited for characterizing early tooth development *in vivo*. Prior work has demonstrated this principle in older children, using MRI to characterize permanent molar eruption patterns in the context of early life stress (11). However, this approach has not been systematically applied to the neonatal primary dentition. The HEALthy Brain and Child Development Study (HBCD), which is acquiring whole-head T2-weighted images from approximately 7,500 newborns across 27 sites, provides an unprecedented opportunity for population-scale characterization of dental development at birth (12).

Here, we develop and validate an automated pipeline for neonatal dental MRI phenotyping in the HBCD cohort. We trained a 3D U-Net to segment the primary dentition from T2-weighted newborn images, extracted spatially localized features along the upper and lower dental arches, and derived a multiparametric estimate of dental age. From these measures, we computed a bias-corrected tooth age gap (TAG) indexing relative dental maturity at birth. We then tested whether TAG was associated with clinical and environmental factors implicated in dental development, including infant sex, race, gestational age at delivery, maternal pre-pregnancy BMI, and prenatal tobacco use. We also tested whether dental maturity at birth prospectively predicted age at first tooth emergence and number of teeth at the 1-year visit. Finally, we evaluated whether automated segmentation and age prediction generalized to an independent cohort. Together, these analyses establish neonatal dental MRI as an objective, scalable approach for measuring primary tooth development before the accumulation of postnatal oral exposures.

## 2. Results

### 2.1 Automated Segmentation of Neonatal Dentition

Semi-automatic reference segmentations of whole dentition took ∼2 minutes per sample to generate (n=100 in total, example output shown in Figure 1, right, see Supplementary Materials for video). Reliability was assessed on a subset of 20 images: intra-rater and inter-rater (JMR) Dice coefficients exceeded 0.98, indicating reproducible manually supervised segmentation.

**Figure 1.**
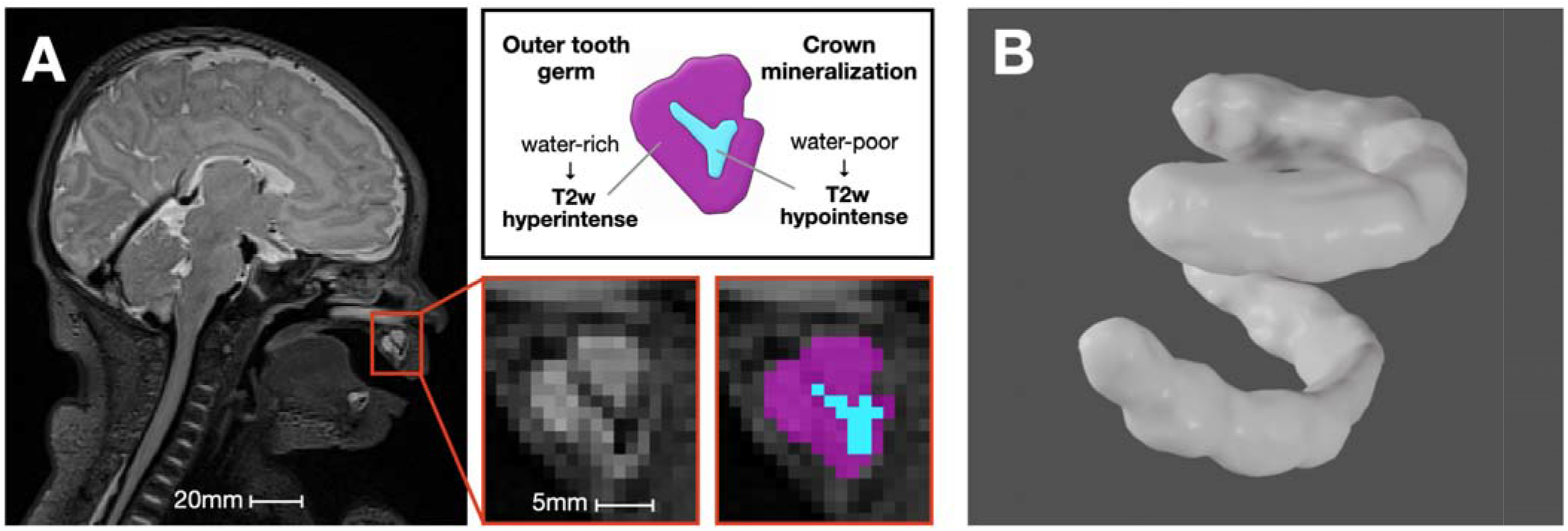
Automated segmentation of neonatal dentition. (A) Representative T2-weighted neonatal MRI showing manual segmentation of an individual tooth (left). The water-rich outer tooth germ appears hyperintense (violet color label) and the mineral-dense inner crown appears hypointense (cyan label) on T2-weighted imaging. (B) A 3D rendered semi-automatic segmentation of the whole upper and lower dentition is shown.

These labeled images were used to train a 3D U-Net under 5-fold cross-validation with online augmentation (5x multiplier per training sample). Training curves demonstrated stable convergence by epoch ∼60, with no evidence of overfitting. The final model (epoch 100) achieved a mean Dice coefficient of 0.94 across held-out folds, with all folds exceeding 0.93. The five trained fold models were applied to all remaining HBCD Release 2.0 scans, and a single consensus segmentation per subject was derived by retaining only voxels labeled as dentition by at least four of five models (4-of-5 supermajority vote). After segmentation quality control, n = 1,545 newborns had usable dentition segmentations; the primary analytic sample was n = 1,433 after excluding scans with missing anthropometric data.

### 2.2 MRI-Based Dental Features Predict Postmenstrual Age

From each consensus segmentation, we extracted 201 image-derived features capturing tooth volume, mineralization, and arch geometry at three hierarchical levels: whole-dentition, per-arch (upper and lower), and along 10 spatially localized segments per arch (Methods 4.4). To derive a summary measure of dental maturity, we used lasso regression (λ_1_⍰ ⍰selection) to predict postmenstrual age (PMA) at MRI from the full set of dental features. Across 100 random 80/20 train-test splits, dental features predicted PMA with a mean R^2^=0.30 and a mean absolute erro (MAE) of 6.7 days (Figure 2A). For comparison, a model using standard anthropometric measures available at birth (birth weight, length, and head circumference) predicted PMA with mean R^2^ = 0.22 using identical methods in the same sample (Figure 2B). Dental features significantly outperformed anthropometric measures (paired t-test across 100 randomly split iterations, p < 10^-10^). These findings remained qualitatively unchanged when limited to participants born full-term. Among the most consistently selected features across iterations were mandibular arch width and molar-region volume, each of which has a clear anatomical interpretation in the context of dental growth. Applying the HBCD-trained segmentation model and dental-age prediction weights to the external UPSIDE sample yielded R^2^ = 0.27 (MAE = 14 days, Supplementary Figure 1), confirming transfer to data collected independently at a different site under a distinct protocol.

**Figure 2.**
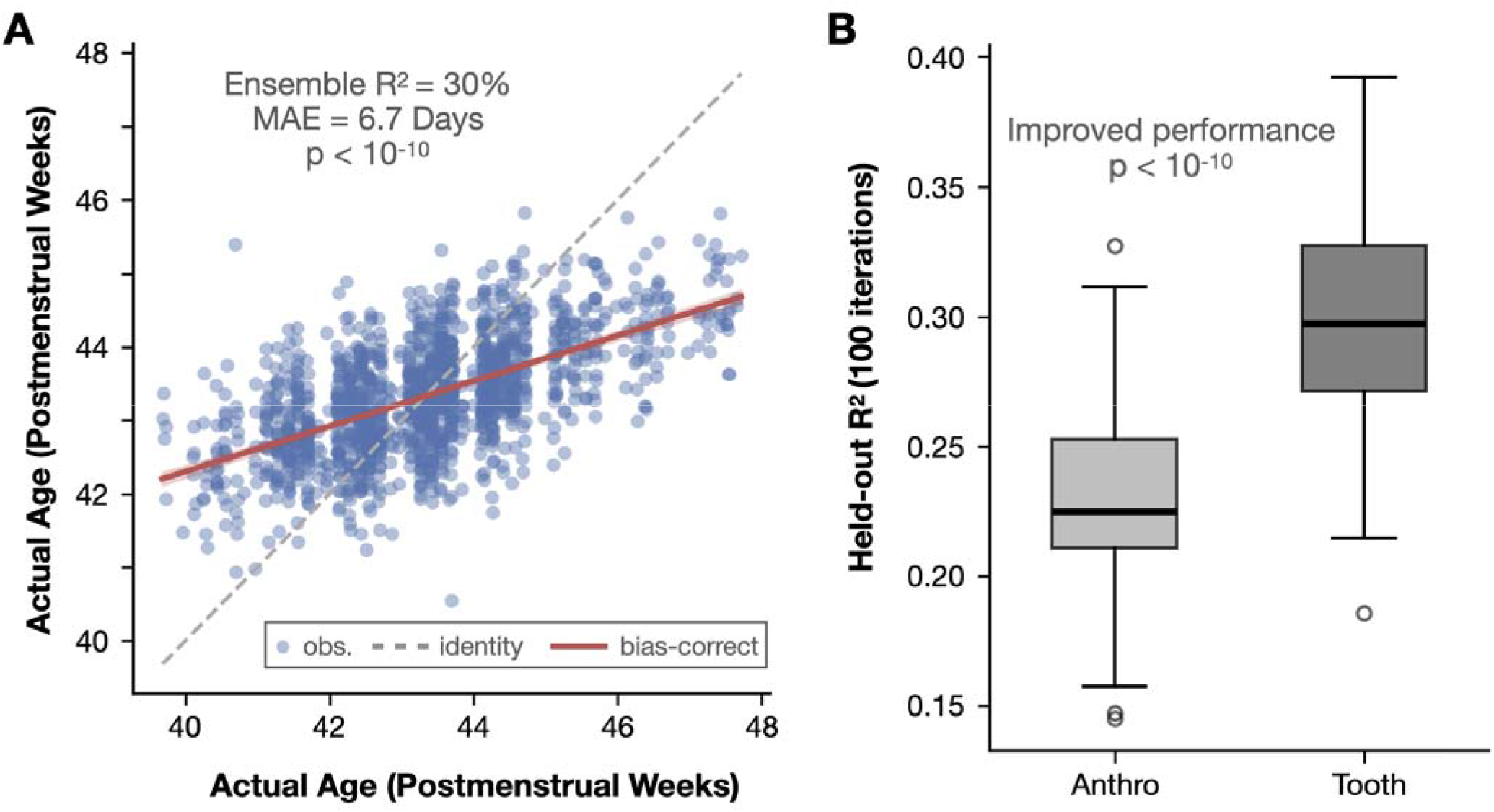
MRI-based dental features predict postmenstrual age and define the tooth age gap. (TAG. (A) Predicted versus actual postmenstrual age (PMA) at MRI for the ensemble lasso model applied to the full sample (n =1,433). The dashed gray line indicates identity prediction. The red line illustrates the bias-correction regression, whose residuals define the tooth age gap (TAG): positive residuals indicate greater-than-expected dental maturity for a given PMA, negative residuals indicate less-than-expected maturity. (B) Distribution of out-of-sample R^2^ across 100 random 80/20 train-test splits for four feature sets: anthropometric measures (birth weight, length, head circumference), and all 201 dental features (regularized). Dental features significantly outperformed anthropometric measures (paired t = 15.3, p < 10^-10^).

### 2.3 Tooth Age Gap Is Associated with Prenatal and Perinatal Factors

We computed a tooth age gap (TAG), defined as the difference between predicted dental age and PMA after correction for regression-to-the-mean bias (Figure 2A, red line), to index each infant’s relative dental maturity at birth. In per-exposure linear mixed-effects models with a random intercept for HBCD site (n = 1,433; 27 sites), TAG was significantly associated with gestational age at delivery (β = −0.034, p < 0.001) and pre-pregnancy BMI (β = 0.008, p < 0.001; approximately 0.035 weeks per 5 kg/m^2^ increase). The unadjusted association between prenatal tobacco use and TAG did not reach significance (β = −0.090, p = 0.092).

In a full model including all exposures, infant sex, race, and image quality covariates (n = 1,326; Figure 3), TAG was associated with infant sex (β_male_ = 0.298, t = 8.75, p < 0.001), with males showing greater dental maturity than females. TAG differed by parent-reported race, with Black infants as the reference group, White infants had lower TAG (β = −0.164, p < 0.001), as did infants in the Other race category (β = −0.177, p < 0.001), corresponding to greater adjusted dental maturity among infants whose parent-reported race was Black. Gestational age at delivery remained significant (β = −0.021, t = −2.17, p = 0.030). However, because PMA reflects both gestational age at delivery and postnatal age at imaging, this association supports a role for birth timing in neonatal dental maturity but does not by itself distinguish prenatal from early postnatal contributions. Pre-pregnancy BMI also remained significant (β = 0.007, t = 3.11, p = 0.002), with higher BMI associated with greater dental maturity. Maternal prenatal tobacco use was associated with reduced dental maturity after covariate adjustment (β = −0.110, t = −2.06, p = 0.039). However, this association was not apparent without covariate adjustment (p = 0.092) and attenuated when head circumference was additionally included (p = 0.096), consistent with shared variation in fetal growth. Image quality covariates were largely non-significant, except for QC PCA component 4 (p < 0.001). Notably, the primary biological associations were unchanged after adjustment for image quality covariates.

**Figure 3.**
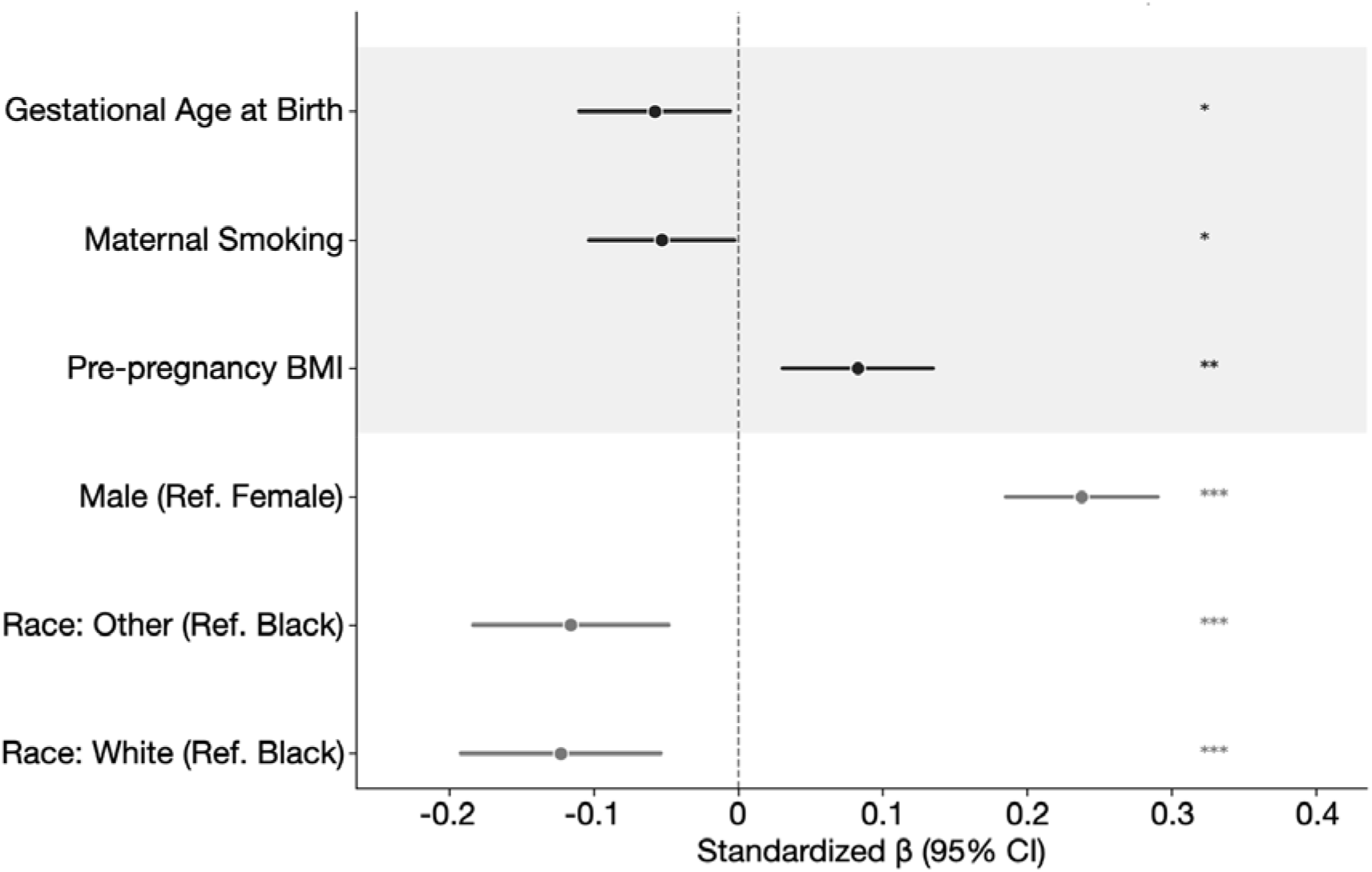
Tooth age gap is associated with gestational exposures. Forest plot of standardized coefficients. (β ±95% CI) from a linear mixed-effects model predicting the tooth age gap from gestational exposures, demogr phic variables, and image quality covariates, with recruitment site as a random intercept (n = 1,326). Males showed greater dental maturity than females (p < 0.001), Black infants showed greater maturity relative to White and Other groups (p < 0.001), and maternal prenatal tobacco use was associated with reduced maturity (p = 0.039). Pre-pregnancy BMI was positively associated with dental maturity (p = 0.002). Image quality covariates were largely non-significant. Vertical dashed line indicates β = 0. *p < 0.05, **p < 0.01, ***p < 0.001.

### 2.4 Dental Maturity at Birth Is Prospectively Associated with Postnatal Eruption

A central question is whether dental maturity assessed at birth carries forward to postnatal eruption outcomes. HBCD Release 2.0 included two parent-reported dental milestones collected at the 9–15-month visit: age at first tooth emergence (ordinal, 6 categories) and number of erupted teeth at approximately 1 year of age (mean age at assessment = 53.0 ± 6.3 weeks). In linear mixed-effects models with a random intercept for site, greater TAG was associated with earlier first tooth emergence (β = −0.307, t = −3.01, p = 0.003, n = 354; Figure 4A) and a greater number of erupted teeth at age 1, adjusting for age at assessment (β = 1.013, t = 4.48, p < 0.001, n = 359; Figure 4B). These associations were robust to adjustment for the full covariate set used in the exposure models (β = −0.300, t = −2.82, p = 0.005, n = 337; β = 0.827, t = 3.49, p < 0.001, n = 342) and to the additional inclusion of head circumference (β = −0.304, p = 0.006; β = 0.814, p < 0.001). Site intraclass correlations were negligible (ICC < 0.013 for both outcomes).

**Figure 4.**
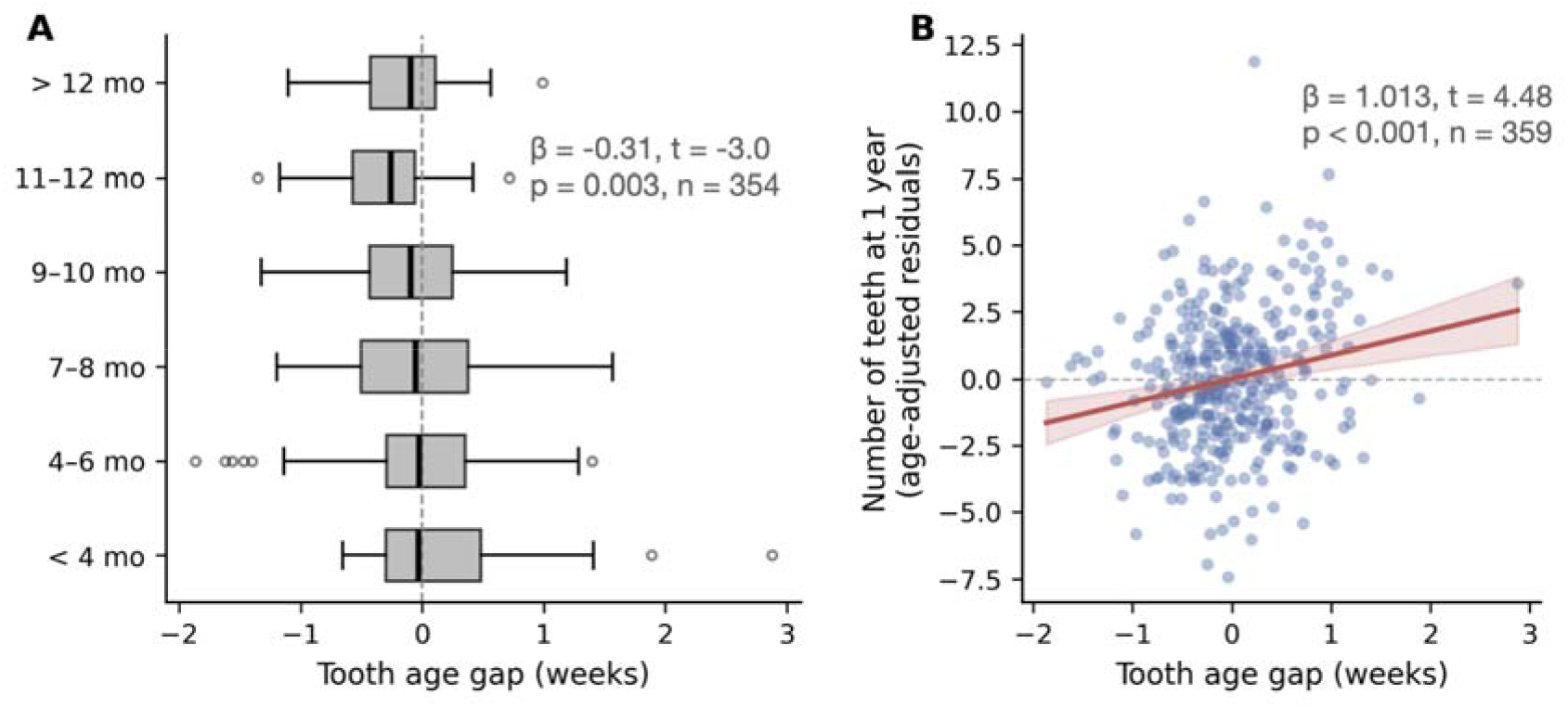
Dental maturity at birth is prospectively associated with postnatal eruption outcomes. (A) Tooth age gap by parent-reported age at first tooth emergence. Boxplots show the median and interquartile range; individual observations are overlaid as jittered points. Greater dental maturity at birth (higher TAG) was associated with earlier first tooth emergence. (B) Tooth age gap versus number of erupted teeth at the approximately 1-year visit, adjusted for child age at assessment. The regression line with 95% confidence band is shown. Greater dental maturity at birth predicted more erupted teeth at age 1.

## 3. Discussion

We demonstrate that neonatal MRI can be used for automated, scalable, in vivo assessment of pre-/perinatal dental mineralization and morphology. Because ionizing-radiation methods are generally not ethically permissible for pediatric research, MRI uniquely enables population-scale study of dental development at the prenatal-postnatal transition. We developed a segmentation and feature extraction pipeline that captures regional variation in tooth volume, mineralization status, and arch geometry, and from these measures derived an individualized index of dental maturity at birth. MRI-derived dental features: 1) substantially outperformed conventional anthropometric measures in predicting postmenstrual age, 2) were associated with demographic, prenatal, and perinatal factors, including infant sex, race, gestational age, pre-pregnancy BMI, and maternal tobacco use, and 3) were prospectively associated with postnatal eruption milestones in a subsample followed longitudinally. Collectively, these findings position the neonatal dentition as a measurable, prenatal-exposure-sensitive index of the biological embedding of intrauterine experience. This index is derivable from routinely acquired brain MRI, operates at population scale, and connects directly to a developmental process underlying childhood oral health.

Dental age estimation is well established in anthropology, orthodontics, and forensic medicine (13, 14), but its application to newborns has been limited because primary teeth are unerupted and difficult to measure directly. In this study, MRI-derived dental features predicted postmenstrual age more accurately than standard anthropometric measures, indicating that the neonatal dentition contains developmental information not captured by global body size. Importantly, this MRI-derived maturity index was not only age-related but prospectively meaningful in that greater TAG at birth was associated with earlier first tooth emergence and more erupted teeth by age 1. These findings support TAG as an objective in vivo measure of dental maturity at birth, bridging the gap between prenatal tooth development and postnatal eruption outcomes.

TAG was also associated with several demographic, prenatal, and perinatal factors previously implicated in dental development. Male infants showed greater dental maturity at birth, and TAG differed by parent-reported race, with Black infants showing greater maturity than White and Other groups. Although evidence linking sex and race to eruption timing is mixed (7, 15) and may reflect socioeconomic, environmental, and other contextual factors, prior work has reported population-level differences in deciduous tooth size (16) and eruption chronology (15). These differences should not be interpreted as intrinsic biological effects of race, but rather as population-level differences that may reflect genetic ancestry, socioeconomic conditions, environmental exposures, nutrition, healthcare access, and structural factors. In this context, the observed TAG differences suggest that the measure captures systematic developmental variation rather than measurement noise.

Maternal pre-pregnancy BMI was positively associated with TAG, consistent with prior reports linking higher pre-pregnancy BMI to greater infant tooth counts (17), although attenuation after adjustment for birth size in prior work suggests that fetal growth may partly contribute to this association. Conversely, prenatal tobacco use was associated with lower TAG only after covariate adjustment and attenuated after inclusion of head circumference, consistent with the possibility that tobacco-related differences in dental maturity overlap with broader fetal growth restriction.Together, these results support TAG as a prenatal-exposure-sensitive measure of neonatal dental development, while emphasizing that individual exposure associations should be interpreted as construct-validating rather than causal.

A key consideration for the broader impact of this work is the portability of the pipeline to existing newborn MRI datasets. The HBCD-trained segmentation model and age-prediction coefficients were applied without modification to the independent UPSIDE cohort and recovered a significant age-maturity relationship (R^2^ = 0.27). Although reduced relative to the within-HBCD estimate (R^2^ = 0.30), possibly due to differences in scanner intensity and MRI signal dropout due to coil sensitivity near the mandibular arch, this suggests that neither the segmentation model nor the feature weights are grossly overfit to HBCD-specific acquisition characteristics. The large, multisite training sample and regularized regression framework help distinguish stable age-related signal from noise across 201 features, a distinction that smaller cohorts could not support independently. The resulting normative models are thus designed for application to smaller, deeply phenotyped cohorts where exposure data are rich but sample sizes preclude fitting these models independently. The segmentation pipeline, pretrained weights, and ensemble coefficients will be publicly released so that any group with neonatal T2-weighted MRI can derive population-referenced dental maturity estimates without retraining.

Despite the strengths of the current study, several limitations should be considered for the interpretation of our findings. First, the current pipeline approximates individual tooth positions using 10 equal arc-length segments along each arch skeleton, rather than parcellating individual teeth. While sufficient for capturing regional developmental gradients, the current version limits tooth-level sensitivity of the derived features. Individual tooth-level segmentation is feasible but requires substantially more labor-intensive training data; this is planned as a next-generation extension to this work. Second, tissue classification relies on a probabilistic Gaussian Mixture Model applied to T2 intensity, and the boundary between mineralized and unmineralized tissue has not been validated against histological ground truth at the time of MRI. Future validation against histology or growth-mark analyses in exfoliated teeth may help calibrate MRI-derived tissue classifications (6). Third, eruption outcomes were parent-reported at the 9 to 15 month visit, introducing both recall error and quantization into coarse ordinal categories that may underestimate the strength of associations reported here. Finally, the tooth age gap is derived from cross-sectional data at a single neonatal timepoint. Longitudinal validation, tracking dental maturation through subsequent imaging timepoints and linking it to eruption as it occurs, is needed to help further establish whether the TAG captures a stable trait-like measure or a developmental snapshot. Extension of the segmentation pipeline to 6-month and 1-year MRI in future versions will enable this.

### Conclusion

Collectively, this study demonstrates that neonatal brain MRI can reveal a previously inaccessible dimension of human development, the maturity of the unerupted primary dentition at birth. By combining automated segmentation, spatially resolved image features, and normative age prediction, we derive an individualized tooth age gap that captures dental developmental variation beyond global body size, relates to prenatal and perinatal factors, and prospectively predicts postnatal eruption milestones. These findings establish the neonatal dentition as a scalable in vivo readout of developmental variation at the prenatal-postnatal transition. More broadly, they show that population MRI can make hidden developmental systems measurable in living newborns, opening a new window into how intrauterine experience is reflected in early biological development and later health-relevant phenotypes.

## 4. Materials and Methods

### 4.1 Study Cohorts

Data were drawn from two independent cohorts. All procedures were approved by the relevant institutional review boards, and written informed consent was obtained from parents or legal guardians. The primary sample came from the HEALthy Brain and Child Development (HBCD) Study, a multi-site prospective cohort designed to characterize brain and behavioral development from the perinatal period through early childhood (12). HBCD is enrolling approximately 7,500 families across 27 U.S. sites, including a nationally representative sample enriched for prenatal substance use (25%) and socioeconomically matched controls (12). For this study, we used HBCD Release 2.0 (early 2026), which includes neonatal T2-weighted MRI and two parent-reported postnatal dental outcomes: age at first tooth emergence and number of erupted teeth at approximately 1 year of age. The present analytic sample was restricted to infants with usable neonatal T2-weighted MRI covering the dentition, postmenstrual age at MRI <48 weeks, successful dentition segmentation and quality control, and non-missing anthropometric/covariate data required for the primary analyses.

The external validation cohort was drawn from the Understanding Pregnancy Signals and Infant Development (UPSIDE) study, a prospective pregnancy-birth cohort based in Rochester, NY, and part of the NIH ECHO program (for imaging details see Rasmussen et al. (18)). UPSIDE enrolled English-speaking pregnant women at less than 14 weeks gestation, with exclusion criteria including history of psychotic illness, known substance abuse, and major neuroendocrine disorder. The cohort has collected extensive biological, psychological, and health data at each trimester and through postnatal follow-up. Neonatal MRI was acquired using a protocol harmonized with HBCD, as both are derived from the Lifespan Baby Connectome Project acquisition framework (19). For external validation, this cohort was restricted to images with a postmenstrual age range consistent with the HBCD study (<48 weeks).

### 4.2 MRI Acquisition

Both cohorts acquired high-resolution T2-weighted anatomical images as part of whole-head imaging batteries designed for neonatal brain development studies (12, 19). T2-weighted sequences exhibit hyperintensity to tissues with long transverse relaxation times (i.e., free/unbound water), making them particularly well suited for characterizing early tooth development: the water-rich outer tooth germ appears hyperintense while the mineral-dense inner crown appears hypointense, producing natural tissue contrast at spatial dimensions well above the acquisition resolution(20). Acquisition parameters for both cohorts are summarized in Table 1.

**Table 1.**
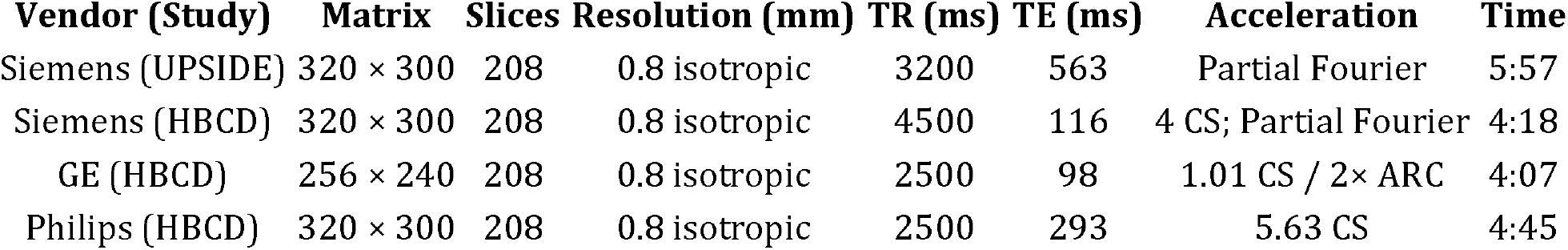
T2-weighted image acquisition parameters. Image acquisition parameters were reasonably harmonized across studies and scanner vendors, including image matrix, slice coverage, spatial resolution, and T2-weighted contrast. Notable differences included variation in TR, TE, acceleration strategy, and the use of compressed sensing in HBCD acquisitions, which enabled reduced scan times.

### 4.3 Dentition Segmentation

One hundred HBCD T2-weighted images were segmented using *nnInteractive*, a 3D promptable segmentation tool that enables rapid semi-automatic labeling via user-defined scribble annotations along the dental arch in a single slice (21). Each segmentation required <5 minutes. All labels were generated by a single trained rater (SS) and adjudicated by an expert in dental anatomy (FBB). Reliability was assessed on a subset of 20 images segmented twice by the primary rater (intra-rater) and once by a second rater (inter-rater, JMR); all Dice coefficients exceeded 0.98, thus demonstrating rater independence.

A 3D U-Net (22) was implemented in PyTorch with three encoder levels and a bottleneck, using a base feature count of 16. Each encoder block comprised two 3×3×3 convolutional layers with InstanceNorm3d and ReLU activation, followed by MaxPool3d(2) for downsampling. The decoder used ConvTranspose3d (kernel size 2, stride 2) for upsampling, with skip connections concatenating encoder and decoder feature maps at each level. The final layer concatenated the output of the last decoder block with the first encoder block’s feature maps (a multi-scale fusion step) before a 1×1×1 convolution with sigmoid activation to produce voxel-wise probability maps.

The model was trained under 5-fold cross-validation with 100 epochs per fold, a batch size of 3, Adam optimizer (learning rate 1×10^−4^), Dice loss, mixed-precision training, and gradient clipping (max norm = 1.0). Each training sample was augmented 5 times using TorchIO RandomAffine transforms (±5° rotation, ±2 voxel translation), yielding an effective training set of 400 image-label pairs per fold. Prior to training, all volumes were min-max normalized and smart-cropped to a fixed target shape of 160×224×256 voxels: the crop was anchored to the superior-most extent of the head (to avoid excluding inferior dental anatomy) and centered on the remaining two axes, with zero-padding as needed.

All five fold models were applied to each unseen image, producing five binary segmentation masks per subject. A consensus segmentation was derived by retaining only voxels classified as dentition by at least four of five models (4-of-5 supermajority vote), favoring specificity over sensitivity.

### 4.4 Feature Extraction

Each consensus segmentation was processed through an automated feature extraction pipeline to derive 201 image-derived features per subject. The segmentation was first parcellated into upper and lower arches by identifying the two largest connected components and assigning them based on their center-of-mass position along the superior-inferior axis. Prior to skeletonization, each arch mask was regularized by filling internal holes, applying Gaussian smoothing (σ = 1.5), and re-thresholding at 0.5 to produce a smooth, topologically simple volume.

Each regularized arch was then skeletonized in 3D to extract a midline tract. Skeleton points were ordered by greedy nearest-neighbor traversal starting from the endpoint with the smallest coordinate along the left-right axis, ensuring consistent orientation across subjects. Each ordered skeleton was divided into 10 equal arc-length segments, roughly corresponding to individual tooth positions along the arch.

Tissue within the segmentation mask was classified into three types. A two-component Gaussian Mixture Model was fit to the T2 intensities of all voxels within the mask, identifying a hyperintense component (water-rich tooth germ) and a hypointense component (mineral-dense crown). A distance-transform-based edge probability function (sigmoid transition with a threshold of 2 voxels from the mask boundary) was combined with the GMM posterior probabilities to define an edge class capturing the transition zone. Final voxel classification was assigned by argmax of composite scores: hyperintense (label 1), hypointense/mineralization (label 2), and edge (label 3).

Volume (mm^3^) and mean T2 intensity were computed for each tissue class at three hierarchical levels: whole dentition (6 features), per arch (12 features), and per segment (10 segments × 2 arches, with 3 tissue volumes, 3 mean intensities, and 3 center-of-mass coordinates per segment, plus arch width and arc length; 183 features). This spatially localized hierarchy captures regional developmental variation along the dental arch.

### 4.5 Quality Control

Five quality control criteria were applied, each assessed independently for the upper and lower arches. Positional outliers were identified by computing the mean squared Euclidean distance of each subject’s segment center coordinates from the population mean and flagging values exceeding ±2 standard deviations. Shape outliers were identified using rigid Procrustes analysis (without scaling or reflection) to compute the disparity between each subject’s arch shape and the population mean shape, again flagging at ±2 SD. Volume outliers were identified by flagging total arch volumes outside ±2 SD. Subjects with missing segment coordinates (indicating failed skeletonization) were also excluded. Finally, subjects with missing postmenstrual age at MRI or PMA ≥ 48 weeks were excluded. The final analytic sample after all QC exclusions comprised 1,433 newborns from 1,545 pre-QC subjects.

Principal component analysis was applied to numeric T2 image quality metrics from MRIQC(23), and the first five components were retained as covariates in downstream models.

Anthropometric measures (birth weight, length and, head circumference) were winsorized at ±3 SD.

### 4.6 Age Prediction and Tooth Age Gap

All 201 dental features were z-scored. Lasso regression (α = 1.0) was used to predict postmenstrual age at MRI, with the regularization parameter selected via the λ_1_⍰ ⍰ rule: from a dense grid of penalty values evaluated under 10-fold cross-validation, we selected the largest penalty whose mean cross-validated error was within one standard error of the minimum; favoring a sparser, more regularized model(24). This procedure was repeated across 100 random 80/20 train-test splits. Model performance was evaluated as the squared Pearson correlation between predicted and actual PMA (R^2^). Four feature sets were compared: anthropometric measures only (birth weight, length, head circumference), PCA-reduced dental features (eigenvalue > 5 and > 1 thresholds), and all 201 dental features. Dental versus anthropometric performance was compared via paired t-test across the 100 split iterations.

An ensemble prediction was derived by averaging the lasso coefficients (intercept and betas) across all 100 iterations and applying them to the full z-scored feature matrix to produce a single predicted dental age per subject (and carried forward for tooth age gap, TAG, definition). A stepwise model was also constructed from the 10 features most frequently selected (non-zero coefficient) across iterations, using BIC as the selection criterion. The tooth age gap was defined as predicted dental age minus PMA. Bias correction was applied by regressing predicted age on actual age and extracting the residuals, which represent deviation from the age-expected prediction, analogous to the “brain age gap” in neuroimaging (25).

### 4.7 Exposure Associations

Associations between the bias-corrected tooth age gap and gestational exposures were tested using linear mixed-effects models with recruitment site as a random intercept. Parsimonious models were first fit for each exposure individually (TAG ∼ exposure + [1|site]) to estimate site-adjusted associations with gestational age at delivery, pre-pregnancy BMI, and maternal prenatal tobacco use (binarized as any vs. none). A full model was then fit including all three exposures simultaneously alongside infant sex, infant race (Black, White, Other), and five T2 image quality PCA components as covariates. A sensitivity analysis additionally included infant head circumference. Race was coded from HBCD demographics following the categories provided by the study, with Black as the reference category. Sex was included in the full model as a covariate to control for demographic confounding of the exposure estimates, but was not treated as an exposure of interest; sex × exposure interaction terms were tested separately. Sample sizes varied across model tiers due to covariate missingness (parsimonious n = 1,433; full n = 1,326).

### 4.8 Eruption Associations

Two parent-reported dental milestones from the HBCD 9–15 month visit were examined as outcomes. Age at first tooth emergence was reported as an ordinal variable with six categories (< 4 months through > 12 months); responses coded as 777 (don’t know) or 999 (declined) were excluded. Number of erupted teeth at approximately 1 year of age was treated as a continuous variable, with the same exclusions. Parsimonious models were fit with the bias-corrected tooth age gap as the primary predictor: for first tooth emergence, TAG + (1|site); for number of teeth, TAG + child age at assessment + (1|site), as age at the visit is an intrinsic determinant of tooth count. Full models additionally adjusted for gestational age at delivery, pre-pregnancy BMI, infant sex, race, maternal tobacco use, and five image quality PCA components, with a further sensitivity analysis including head circumference. Site intraclass correlations were negligible for both outcomes (ICC < 0.013); where full models with random intercepts failed to converge due to small cluster sizes (n ≈ 340 across 27 sites), ordinary least squares regression was used, which yields equivalent estimates when site-level variance is near zero.

### 4.9 External Validation

To assess generalizability, the HBCD-trained segmentation model was applied to UPSIDE neonatal T2-weighted images without retraining, using the same supermajority vote consensus and feature extraction pipeline. The ensemble lasso coefficients derived from HBCD were then applied to the UPSIDE dental features to generate predicted dental age. Age prediction performance was evaluated via R^2^ in the independent sample. This procedure tests the transferability of both the automated segmentation and the learned age-maturity relationship to data collected at a different site under a related but distinct acquisition protocol, without conflating segmentation generalizability with model retraining.

## Acknowledgements

We thank the participant volunteers for donating their time to the HBCD and UPSIDE projects and the generosity of the National Institutes of Health for making the data publicly available.

## Funding

NICHD R00 HD-100593 to JMR; NIMH R01 MH-138481 to JMR; NIH grant # OD023349 to TO’C

## Authors’ contributions

YM, FBB, TGO, JX, JMR conceptualized and designed the study. JMR developed the segmentation and analysis pipeline, trained and validated the neural network, performed all statistical analyses, and co-wrote the manuscript. YM and JX compiled and curated UPSIDE cohort data, contributed to data analysis, and co-wrote the manuscript. TGO obtained funding for and directed the UPSIDE cohort and co-wrote the manuscript. FBB provided expert adjudication of dental segmentations and co-wrote the manuscript. YM, FBB, JX contributed domain expertise in dental anatomy. SS performed manual segmentation of the training dataset. All authors reviewed and approved the final manuscript.

## Conflicts of Interest Statement

All authors have no conflicts of interest to declare.

## Prior Presentation

This work has not been previously presented.

## Data, Materials, and Software Availability

Underlying individual participant data and corresponding data dictionaries are shared as part of the HBCD data repository at the NIH Brain Development Cohorts (NBDC) Data Sharing Platform (https://www.nbdc-datahub.org/). All variables used in this manuscript are denoted by their data dictionary name. Note that future HBCD data release variable names may not be consistent with those used here. All derived data (e.g., segmentations and image derived tooth measures) can be recreated through public access code (https://github.com/jerodras/hbcd_teeth), and pretrained/derived outputs deposited or available at a stable repository, with restricted HBCD data through NBDC DUAs.

---

**Supplementary Figure 1.**
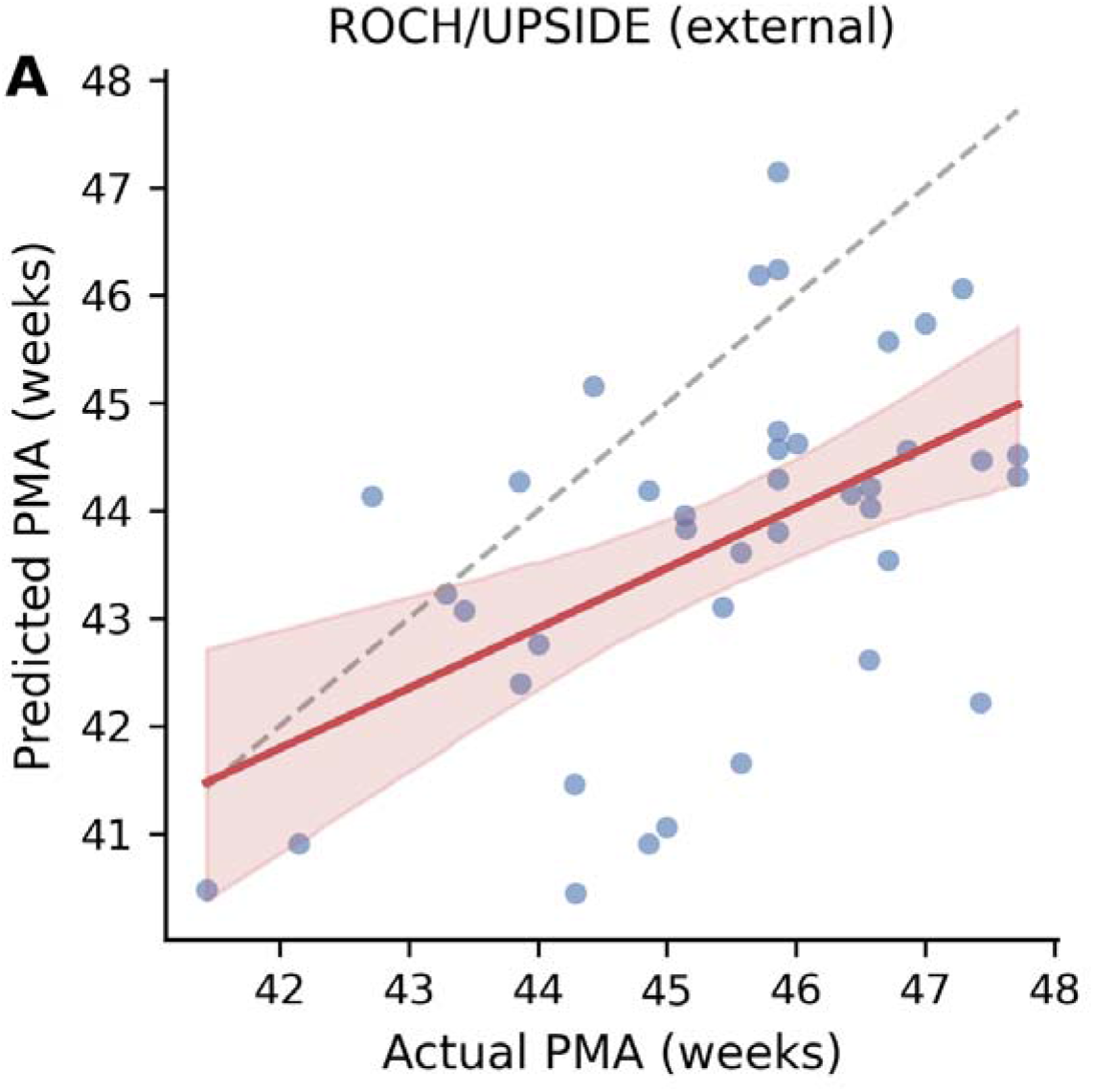
External validation in the UPSIDE/ROCH cohort. Predicted versus actual postmenstrual age for the HBCD-trained ensemble model applied to the independent UPSIDE/ROCH cohort (n = 40, restricted to PMA ≤ 48 weeks to match the HBCD training range). Dashed gray line indicates identity. R^2^ = 0.27, MAE = 14 days.

**Supplementary Figure 2.**
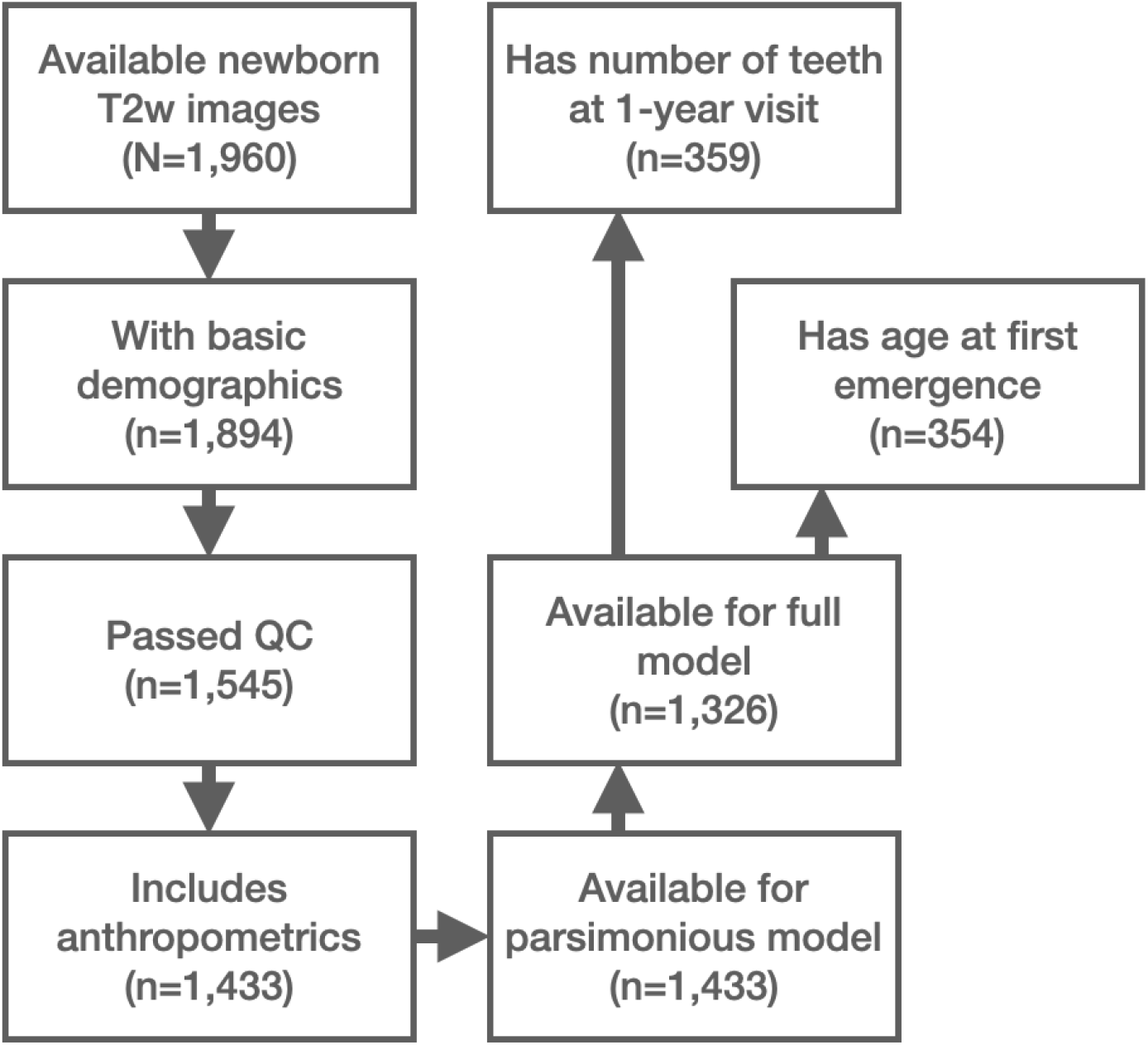
CONSORT figure.

## Notes

### Competing Interest Statement

The authors have declared no competing interest.

## References

1. C. Centers for Disease, Prevention (2024) Oral Health Surveillance Report: Dental Caries, Tooth Retention, and Edentulism, United States, 2017–March 2020. (U.S. Department of Health and Human Services, Atlanta, GA).

2. F. S. Costa et al., Developmental defects of enamel and dental caries in the primary dentition: A systematic review and meta-analysis. Journal of Dentistry 60, 1–7 (2017).

3. G. Ntani et al., Maternal and early life factors of tooth emergence patterns and number of teeth at 1 and 2 years of age. Journal of Developmental Origins of Health and Disease 6, 299–307 (2015).

4. J. J. Warren et al., Tooth Eruption and Early Childhood Caries: A Multisite Longitudinal Study. Pediatric Dentistry 43, 287–289 (2021).

5. D. S. Koussoulakou, L. H. Margaritis, S. L. Koussoulakos, A curriculum vitae of teeth: evolution, generation, regeneration. International Journal of Biological Sciences 5, 226–243 (2009).

6. J. Estivals et al., Insight into perinatal health by investigating the neonatal line: A systematic review. Archives of Oral Biology 177, 106335 (2025).

7. A. I. Georgiadou, A. Ritsas, A. Arhakis, The Impact of Maternal, Perinatal, and Early Infancy Period on the Eruption Timing of the First Primary Tooth. European Journal of Dental and Oral Health 2, 28–33 (2021).

8. S. G. Reed, S. Fan, C. L. Wagner, A. B. Lawson, Predictors of Developmental Defects of Enamel in Primary Maxillary Central Incisors Using Bayesian Model Selection. Caries Research 58, 30–38 (2024).

9. G. Tapalaga et al., The Impact of Prenatal Vitamin D on Enamel Defects and Tooth Erosion: A Systematic Review. Nutrients 15, 3863 (2023).

10. A. A. Akinkugbe, Does the Trimester of Smoking Matter in the Association between Prenatal Smoking and the Risk of Early Childhood Caries? Caries Research 55, 114–121 (2021).

11. C. L. McDermott et al., Early life stress is associated with earlier emergence of permanent molars. Proceedings of the National Academy of Sciences of the United States of America 118 (2021).

12. C. A. Nelson, J. Frankeberger, C. D. Chambers, An introduction to the HEALthy Brain and Child Development Study (HBCD) study. Developmental Cognitive Neuroscience 69, 101441 (2024).

13. V. Marconi et al., Validity of age estimation methods and reproducibility of bone/dental maturity indices for chronological age estimation: a systematic review and meta-analysis of validation studies. Scientific Reports 12, 15607 (2022).

14. D. H. Ubelaker, H. Khosrowshahi, Estimation of age in forensic anthropology: historical perspective and recent methodological advances. Forensic Sciences Research 4, 1–9 (2019).

15. J. J. Warren et al., Timing of primary tooth emergence among U.S. racial and ethnic groups. Journal of Public Health Dentistry 76, 259–262 (2016).

16. E. F. Harris, B. D. Barcroft, American black-white differences in primary tooth crown dimensions. Dental Anthropology 15, 1–5 (2001).

17. G. Ntani et al., Maternal and early life factors of tooth emergence patterns and number of teeth at 1 and 2 years of age. Journal of Developmental Origins of Health and Disease 6, 299–307 (2015).

18. J. M. Rasmussen et al., Segmenting hypothalamic subunits in human newborn magnetic resonance imaging data. Human Brain Mapping 45, e26582 (2024).

19. B. R. Howell et al., The UNC/UMN Baby Connectome Project (BCP): An overview of the study design and protocol development. NeuroImage 185, 891–905 (2019).

20. C. L. McDermott et al., Early life stress is associated with earlier emergence of permanent molars. Proceedings of the National Academy of Sciences of the United States of America 118, e2105304118 (2021).

21. F. Isensee et al., nnInteractive: Redefining 3D Promptable Segmentation. arXiv, arXiv:2503.08373 (2025).

22. Ö. Çiçek, A. Abdulkadir, S. S. Lienkamp, T. Brox, O. Ronneberger (2016) 3D U-Net: Learning Dense Volumetric Segmentation from Sparse Annotation. in Medical Image Computing and Computer-Assisted Intervention – MICCAI 2016, pp 424–432.

23. O. Esteban et al., MRIQC: Advancing the automatic prediction of image quality in MRI from unseen sites. PLOS ONE 12, e0184661 (2017).

24. J. H. Friedman, T. Hastie, R. Tibshirani, Regularization Paths for Generalized Linear Models via Coordinate Descent. Journal of Statistical Software 33, 1–22 (2010).

25. S. M. Smith, D. Vidaurre, F. Alfaro-Almagro, T. E. Nichols, K. L. Miller, Estimation of brain age delta from brain imaging. NeuroImage 200, 528–539 (2019).

